# When Grammatical Gender Shapes Gender Stereotypes: Neural Evidence for Cross-Linguistic Modulation in Spanish-English Bilinguals

**DOI:** 10.64898/2026.08.25.746218

**Authors:** Francesca Pesciarelli, Maria Christina Huerta-Avila, Jacklyn Jardel, Katherine J. Midgley, Phillip J. Holcomb

## Abstract

Can grammatical gender in a bilingual’s first language shape gender-stereotype processing in a second language? Spanish (L1)-English (L2) bilinguals (n = 28) and English monolinguals (n = 28) completed an event-related potential (ERP) priming task in which English pronouns (SHE/HE) followed gender-stereotyped English nouns, half of which had gender-marked Spanish translation equivalents (e.g., NURSE “enfermera/o”, SURGEON “cirujana/o”), and half unmarked translation equivalents (e.g., SINGER “cantante”, JANITOR “conserje”). Both groups showed asymmetric stereotype priming: male pronouns elicited a larger N400 for incongruent than congruent primes, whereas female pronouns elicited a larger P300 for incongruent than congruent primes. Crucially, only bilinguals showed modulation by Spanish grammatical gender marking: the N400 effect for male pronouns was larger for primes with gender-marked than unmarked Spanish translations. These findings provide neural evidence that grammatical gender in a bilingual’s first language can influence gender-stereotype processing in a second language, linking cross-linguistic activation to social cognition.

## Introduction

The present study examines whether gender stereotype processing in bilinguals is driven exclusively by conceptual-semantic expectations or can also be modulated by grammatical-gender information from the non-target language.

Language comprehension relies on multiple sources of information that allow comprehenders to anticipate upcoming input. One particularly relevant source is gender information, which may be conveyed both by socially grounded stereotypes and by grammatical structure. Role nouns such as *nurse, surgeon*, or *singer* often evoke probabilistic expectations about the likely gender of the referent, reflecting gender stereotypes, that is, socially shared beliefs about the roles, attributes, or occupations more typically associated with one gender than another. Such stereotype-based expectations can influence the processing of subsequent pronouns and other referring expressions (e.g., Banaji & Hardin, 1996; Casado et al., 2023; Ellemers, 2017; Fabre et al., 2015, 2016; Garnham et al., 2002; Pesciarelli et al., 2019; Proverbio et al., 2018; Serafini & Pesciarelli, 2025; Siyanova-Chanturia et al., 2012, 2015) and can do so rapidly, even without conscious awareness and in the absence of explicit gender marking (Pesciarelli et al., 2019). Gender-stereotype processing, therefore, provides a useful domain for investigating how lexical meaning, world knowledge, and linguistic form interact during comprehension.

In event-related potential (ERP) studies, gender stereotype-incongruent continuations have often been associated with modulations in components linked to expectancy, semantic integration, and updating, most notably the N400 and later positivities. The N400, which peaks around 400 ms after stimulus onset and is typically associated with semantic incongruity (Kutas & Federmeier, 2011), is one of the most consistent correlates of stereotype violation, emerging in response to stereotype-incongruent pronouns, faces, sentences, and voices (e.g., Grant et al., 2020; Molinaro et al., 2016; Pesciarelli et al., 2019; Proverbio et al., 2018; Rodríguez-Gómez et al., 2020; Serafini & Pesciarelli, 2025, 2026; Siyanova-Chanturia et al., 2012; Van Berkum et al., 2008). For example, using a noun-pronoun priming paradigm, Pesciarelli et al. (2019) showed that stereotypically gendered nouns influence the processing of subsequent gendered pronouns, with stereotype-incongruent pronouns (e.g., *insegnante-LUI*, “teacher—He”) eliciting a larger N400 than congruent pronouns (e.g., *ingegnere-LUI*, “engineer—HE”).

By contrast, later positive components have yielded a more variable pattern. The P600, which peaks between 500-900 ms, has often been interpreted as reflecting syntactic reanalysis or revision processes triggered by stereotype violation (Canal et al., 2015; Irmen et al., 2010; Lattner & Friederici, 2003; Osterhout et al., 1997; Osterhout & Holcomb, 1996; Proverbio et al., 2017; Su et al., 2016), whereas related positivities such as the P300 and the Late Positive Potential (LPP), associated with context updating and stimulus salience (Donchin, 1981; Hajcak et al., 2010), have shown less consistent involvement. In particular, such later effects have not been reliably observed for incongruent pronouns or words, but have been reported for stereotype-incongruent faces in mixed word-face paradigms (Rodríguez-Gómez et al., 2020; Serafini & Pesciarelli, 2025, 2026).

A growing body of ERP evidence suggests that male and female stereotype violations are not processed symmetrically, with different neural responses emerging depending on the direction of the violation. When a male agent is paired with a female-stereotyped role (e.g., *caregiver-HE*/male face), studies have predominantly reported N400 effects (Proverbio et al., 2018; Serafini & Pesciarelli, 2025, 2026; Siyanova-Chanturia et al., 2012). In contrast, pairing a female agent with a male-stereotyped role (e.g., *carpenter-SHE*/female face) has more often produced later positivities, including the P300, P600, and LPP (Irmen et al., 2010; Proverbio et al., 2017; Serafini & Pesciarelli, 2025, 2026; Su et al., 2016). However, the basis of this emerging asymmetry remains unclear and may involve broader social-cognitive factors, such as the relative markedness of masculine and feminine categories, differences in the representation of male-typed and female-typed roles, or asymmetric baseline expectations about men and women in stereotyped occupations (Eagly & Wood, 2012). For a more detailed discussion of this asymmetry, see Serafini & Pesciarelli (2026).

While the above literature has primarily focused on how gender stereotypes shape online comprehension in monolingual settings, much less is known about how these processes unfold in bilingual speakers. This is an important and still underexplored question, because in bilingual language processing, the target language is not necessarily the only source of information available online. A substantial body of behavioral and ERP evidence suggests that, even when a task is carried out in one language, lexical representations from the non-target language may also become activated (Kroll & Stewart, 1994; Midgley et al., 2008; Sunderman & Kroll, 2006; Thierry & Wu, 2007). For example, Thierry and Wu (2007) showed that Chinese-English bilinguals were sensitive to hidden translation relationships in English word pairs, suggesting automatic activation of L1 lexical representations during L2 processing. Such cross-linguistic co-activation makes it plausible that information encoded in a bilingual’s first language may shape interpretation in a second language, even when it is not overtly expressed in the language of the task. This issue is particularly relevant for gender processing, where bilinguals may rely not only on stereotype-based information associated with role nouns in the target language, but also on grammatical-gender information encoded in the non-target language. Bilingualism, therefore, offers a valuable testing ground for investigating whether gender stereotype processing is driven exclusively by conceptual-semantic expectations or can also be modulated by cross-linguistic co-activation of grammatical gender.

Such cross-linguistic differences become especially salient when bilinguals’ two languages differ in the extent to which gender is morphologically encoded. Spanish-English bilinguals provide a particularly informative case in this respect, because Spanish encodes grammatical gender overtly, whereas English does not. Importantly, Spanish human-denoting nouns also vary in whether gender is morphologically specified: some have distinct masculine and feminine forms (e.g., NURSE “enfermera/o”, SURGEON “cirujana/o”), whereas others are morphologically unmarked or common-gender forms (e.g., SINGER “cantante”, JANITOR “conserje”). If bilingual lexical access involves cross-linguistic co-activation, then English nouns may activate not only stereotype-based gender expectations, but also the grammatical-gender information associated with their Spanish translation equivalents. This distinction between gender-marked and unmarked Spanish translation equivalents makes it possible to test whether grammatical gender in bilinguals’ L1 modulates online gender stereotype processing during L2 English comprehension.

Despite the theoretical and social relevance of this question, empirical evidence on gender stereotype processing in bilinguals remains limited. Available behavioral findings suggest that such processing may vary across languages and cultural contexts (Sato et al., 2013, 2016). Far less is known, however, about its neural underpinnings. To our knowledge, only two ERP studies have examined gender stereotype processing in bilinguals to date. Jankowiak et al. (2025) found reduced stereotype effects during foreign language sentence processing in Polish L1-English L2 bilinguals, suggesting attenuated stereotype activation in L2. By contrast, in a noun-pronoun priming paradigm similar to that of Pesciarelli et al. (2019), Casado et al. (2026) reported a larger gender congruency effect for stereotypical than for neutral primes in bilinguals, whereas no such modulation emerged in monolinguals. In that study, we interpreted this finding as reflecting greater reliance on semantic information during bilingual processing of stereotype-related words. However, despite these initial findings, it remains unknown whether language-specific structural properties, such as grammatical gender, shape the online processing of stereotype-based gender expectations in bilinguals.

The present study investigated whether Spanish grammatical gender modulates the processing of English gender stereotypes in highly proficient Spanish (L1)-English (L2) bilinguals. Using an ERP noun-pronoun priming paradigm adapted from Pesciarelli et al. (2019), we presented English target pronouns (*SHE, HE*) following stereotypically gendered English nouns whose Spanish translation equivalents were either gender-marked (e.g., NURSE “enfermera/o”, SURGEON “cirujana/o”) or unmarked (e.g., SINGER “cantante”, JANITOR “conserje”). This design allowed us to test whether stereotype-based priming in English is further modulated by grammatical-gender information from Spanish. A monolingual English control group was tested on the same materials and with the same paradigm to determine whether any such modulation reflects bilingual cross-linguistic influence rather than a general property of stereotype processing or the experimental materials and task. We expected stereotype-congruent primes to facilitate pronoun processing in both groups, but predicted a stronger priming effect for gender-marked than unmarked translation equivalents only in bilinguals. In line with the increasingly consistent evidence in the literature, we also expected male and female stereotype violations to elicit asymmetric ERP responses in both groups, reflecting a general property of gender stereotype processing rather than bilingual cross-linguistic influence.

## Method

### Participants

Two groups of participants were recruited from San Diego State University via SONA and from the San Diego area through flyers and the NeuroCognition lab participant database. Initially, 29 fluent Spanish-English bilinguals (21 women, 7 men, and 1 non-binary; age range = 18–40, M = 24.59, SD = 5.65) and 28 English monolinguals (18 women, 9 men, 1 preferred not to report; age range = 18–40, M = 24.89, SD = 5.12) were enrolled. One bilingual participant was excluded after reporting exclusive English use at home (100%) on the Bilingual Language Profile (BLP; Birdsong et al., 2012) and showing low Spanish proficiency on the Multilingual Naming Test (MINT; MINTSprint 2.0; Gollan et al. 2023; Spanish score = 33), resulting in a final sample of 28 bilingual participants (21 women, age range = 18–40 years, M = 24.5, SD = 5.73). All participants were right-handed, had normal or corrected-to-normal vision, and reported no history of neurological disorders.

Monolingual participants were highly proficient only in English and had been exposed to English from birth.

Bilingual participants completed the BLP, which assesses Spanish and English language history, including proficiency, exposure, use, and age of acquisition. All participants had been exposed to Spanish from birth. English acquisition began at birth or later (range = 0–17 years, M = 5.14, SD = 4.11). Self-reported proficiency ratings indicated that participants perceived themselves as proficient in both L1 Spanish and L2 English at the time of testing. Language dominance was indexed using the BLP dominance scores, computed as the difference between the two language scores (range 1–250 per language), with negative values indicating Spanish dominance and positive values indicating English dominance. Mean scores were 158.0 (SD = 29.82) for English and 161.9 (SD = 18.81) for Spanish, yielding a dominance score of -3.9, which was not statistically significant, t (24) = -0.39, p = .70 (n = 25; 3 missing BLP dominance scores).

To provide an objective measure of language proficiency, participants completed the MINT (MINTSprint 2.0; Gollan et al., 2024). Bilingual participants completed the task in both Spanish and English, whereas monolingual participants completed the English version only. Accuracy was quantified as the total number of correctly named items (maximum = 80). Bilinguals obtained a mean MINT score of M = 63.50 (SD = 9.69) in English and M = 58.86 (SD = 9.73) in Spanish; a paired-samples t-test indicated that performance in Spanish and English did not differ significantly, t (27) = 1.74, p = .09. Monolinguals’ English MINT performance was M = 76.54 (SD = 2.86). An independent-samples t-test comparing English MINT scores between bilinguals and monolinguals indicated higher scores in the monolingual group, t (54) = -6.83, p < .001.

Participants volunteered and provided written informed consent in accordance with the San Diego State University Institutional Review Board. They were compensated for their time.

### Stimuli

Stimuli consisted of 156 stereotypically gendered English word primes (nouns and adjectives) referring to occupations, social roles, and personal characteristics, selected on the basis of rating questionnaires. Seventy-eight words were associated with male stereotypes (e.g., SURGEON) and 78 with female stereotypes (e.g., NURSE) (see Supplementary Material for the prime-word norming study and full stimulus list). For each stereotype category, half of the words had gender-marked Spanish translations (e.g., NURSE “enfermera/o”, SURGEON “cirujana/o”), and half had gender-unmarked translations (e.g., SINGER “cantante”, JANITOR “conserje”). The four stimulus sets (gender-marked vs gender-unmarked x male-vs. female-stereotypes) were matched on stereotype strength, word frequency (NIM; Guasch et al., 2013), word length in characters, cognate status, and valence (see Table 1 in the Supplementary Material). In addition, 40 filler nouns with no associated gender stereotype (e.g., PARTNER, TRAVELER) were included in each block to reduce the salience of the gender-stereotype words and minimize strategy use. Each prime was followed by a third-person singular pronoun (HE or SHE). Primes were paired with a pronoun to create stereotype-congruent pairs (e.g., marked: SURGEON– HE, NURSE– SHE; unmarked: JANITOR– HE, SINGER– SHE) and stereotype-incongruent pairs (e.g., marked: SURGEON – SHE, NURSE – HE; unmarked: JANITOR – SHE, SINGER - HE). The experiment consisted of two blocks, each comprising 196 trials (39 prime-target trials in each of the four experimental conditions plus 40 filler trials). For each prime, the target pronoun was feminine (SHE) in one block and masculine (HE) in the other. ERP analyses focused on the target pronouns, which constituted the critical stimuli for all comparisons. Thus, each participant completed a total of 392 trials. Prime-target pairs were randomized before presentation. Prior to the main experiment, participants completed a brief practice session consisting of 16 trials using stimuli not included in the experimental set (8 marked and 8 unmarked).

### Design and Procedure

Participants were seated comfortably in a dimly lit, sound-attenuated room. An example of the stimulus presentation sequence is shown in Figure 1. All stimuli (4–13 letter strings) were presented centrally on a computer monitor in white uppercase Arial font against a black background. Each trial began with a central fixation cross (+) displayed for 800 ms, followed by a 500 ms blank screen. The prime word was then presented for 300 ms at the same location, followed by a 200 ms blank interval. Subsequently, the target pronoun (SHE or HE) appeared for 300 ms. The stimulus was followed by an 800 ms blank screen and a final fixation cross, which remained on the screen until the participant pressed a button on a gamepad resting in their lap. The next trial then began. Participants were instructed to indicate, as quickly and accurately as possible, whether the pronoun was female or male by pressing one of two buttons, which were counterbalanced (left and right) across participants. They were asked to remain still and to avoid blinking during stimulus presentation but were allowed to blink and rest during the inter-trial interval while the final fixation cross was displayed.

**Figure 1.**
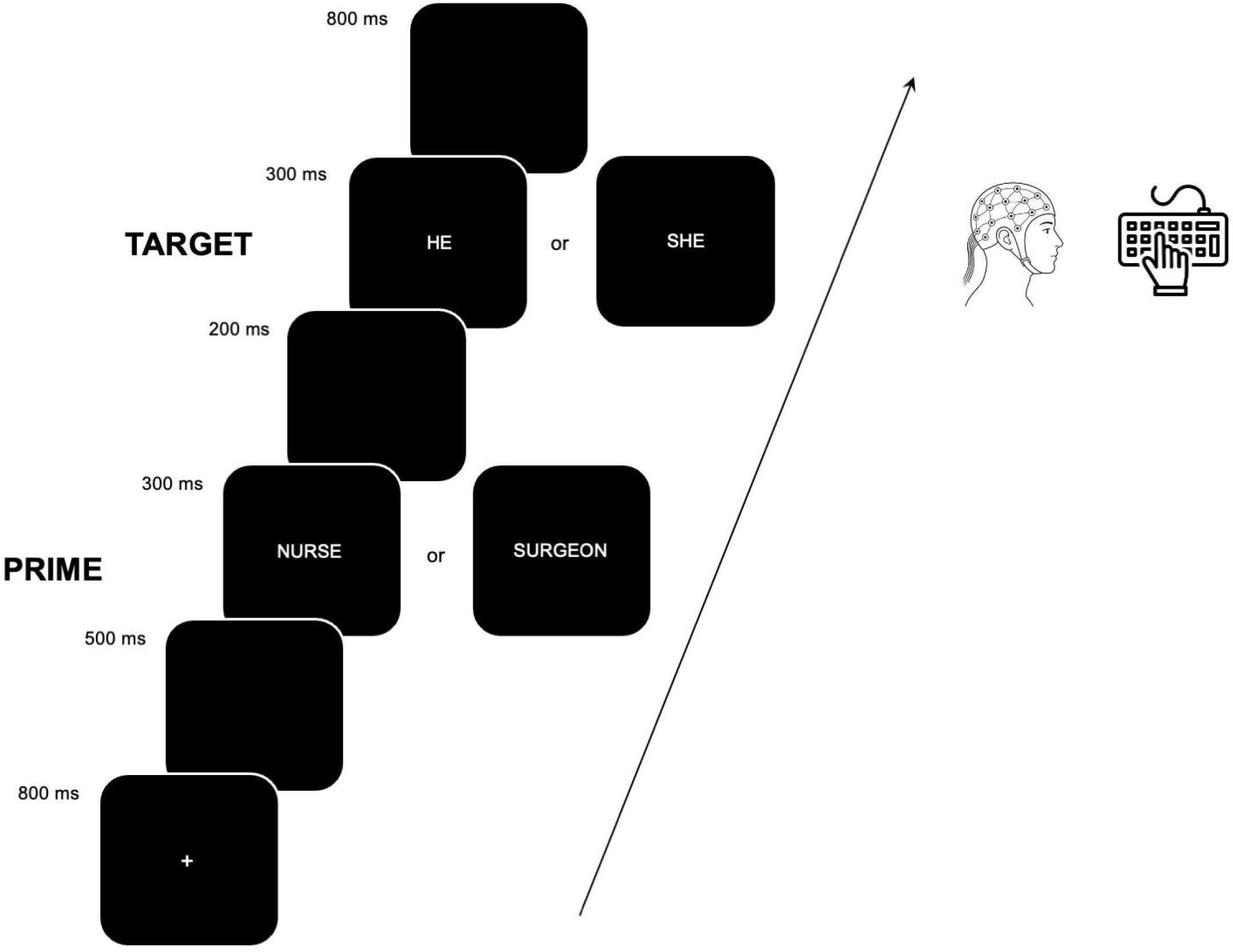
Schematic representation of the experimental paradigm. Each trial began with a fixation cross, followed by a prime word referring to a stereotypically male or female role (e.g., *surgeon, nurse*). The prime was then followed by a target pronoun (*he* or *she*). Prime– target pairs could be congruent or incongruent with respect to gender stereotypes.

### EEG recording and analysis

EEG was continuously recorded from a 32-channel tin electrode cap (Electro-cap International Inc., Eaton, OH). The signal was amplified using a SynAmpsRT amplifier (Neuroscan-Compumedics, Charlotte, NC), with a bandpass of DC-200 Hz and a sampling rate of 500 Hz. In addition, two electrodes were placed below the left eye and at the outer canthus of the right eye to monitor ocular activity. The reference electrode was positioned on the left mastoid, with an additional electrode on the right mastoid to monitor differential mastoid activity. Electrode impedances were maintained below 2.5 kΩ. Data preprocessing was conducted in MATLAB using EEGLAB (Delorme & Makeig, 2004) and ERPLAB (Lopez-Calderon & Luck, 2014). The continuous EEG was band-pass filtered offline using a noncausal Butterworth (0.1–30 Hz; 12 dB/octave roll-off). To facilitate ocular artefact correction, bipolar horizontal (HEOG; horizontal right minus F7 channels) and vertical (VEOG; FP1 minus lower left eye channels) EOG channels were computed. Segments containing extreme voltage values were first excluded from the continuous EEG. Independent component analysis (ICA) was then performed on the original EEG channels. Components reflecting eye blinks were identified based on their temporal and spatial characteristics and removed from the data, after which the EEG signal was reconstructed. The data were then re-referenced offline to the average activity of the two mastoids and segmented into epochs from ™100 to 800 ms relative to target onset, with 100-ms pre-stimulus baseline correction. Artifact rejection was subsequently applied across all channels, rejecting trials with extreme values in any channel. Trials containing artifacts were then removed. Epochs corresponding to correct responses were averaged across eight conditions (mean number of epochs per condition = 37.53, SD = 1.50, range = 32–39). Data loss due to artifacts amounted to 0.60% (SD = 1.21%). For analyses time-locked to target pronoun onset, ERP components were identified based on visual inspection of grand-averaged waveforms and in accordance with previous literature (e.g., Pesciarelli et al., 2019; Serafini & Pesciarelli, 2025, 2026; Siyanova-Chanturia et al. 2012). Two components were identified at frontal (F3, Fz, F4), central (C3, Cz, C4), and parietal (P3, Pz, P4) electrodes within the following time windows: N400 (250-400 ms); P300 (380-550 ms).

### Statistical analysis

Statistical analyses were conducted in JASP (Version 0.95.4). Behavioral analyses were performed on response times (RTs) between 200 ms and 1200 ms, corresponding to 96.54% of trials, in order to exclude anticipatory and delayed responses. Accuracy was not analyzed due to ceiling effects (96%–98% correct across conditions). Both RT and ERP analyses were restricted to trials with correct responses. RTs and ERP mean amplitude were analyzed using repeated-measure ANOVAs with Prime Type (marked, unmarked), Target Gender (female, male), and Congruency (congruent, incongruent) as within-subject factors, and Group (monolinguals, bilinguals) as a between-subject factor. For ERP analyses, Longitude (anterior, central, posterior) and Latitude (left, midline, right) were included as additional within-subject factors. The levels corresponded to the mean activity of F3, Fz, F4 (Anterior), C3, Cz, C4 (Central), P3, Pz, P4 (Posterior), F3, C3, P3 (Left), Fz, Cz, Pz (Midline), and F4, C4, P4 (Right). When Group and/or Target Gender significantly interacted with Congruency, indicating that the priming effect varied by group and/or target gender (the latter being consistent with our previous findings), follow-up analyses were conducted. Specifically, significant two-way interactions were probed using simple effects analyses within the same model, and higher-order interactions (three- and four-way) were examined by conducting separate ANOVAs for monolingual and bilingual participants, and for female and male targets. All post hoc mean comparisons were corrected using the Holm procedure (p < . 05). Degrees of freedom were corrected using the Greenhouse–Geisser method, and only corrected p-values are reported. Partial eta squared (η^2^p) is reported as a measure of effect size. The significance threshold was set at p = .05. Main effects and interactions that did not involve the priming manipulation (i.e., the Congruency factor) were not central to the research question and are therefore not reported.

## Results

### Behavioral results

The omnibus ANOVA on RTs showed a significant main effect of Congruency, F(1, 54) = 19.37, p < .001, η^2^p = .26, with faster responses to pronouns preceded by stereotypically congruent compared to stereotypically incongruent primes. This pattern indicates a reliable priming effect that was consistent across both monolingual and bilingual participants. No other effect of interest reached significance (all ps > .1).

### ERP results

Grand-averaged ERPs elicited by the different experimental conditions are represented in Figure 2.

**Figure 2.**
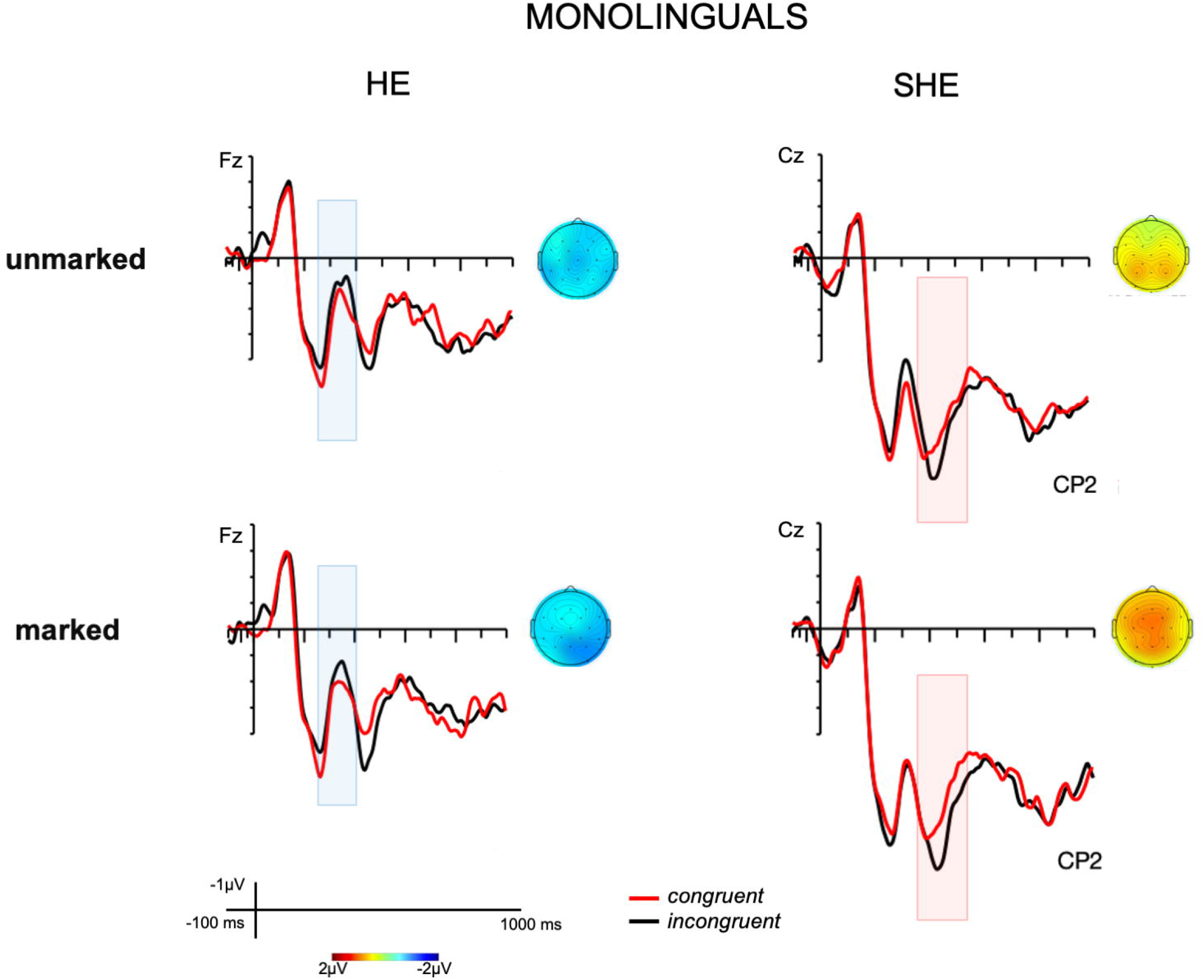
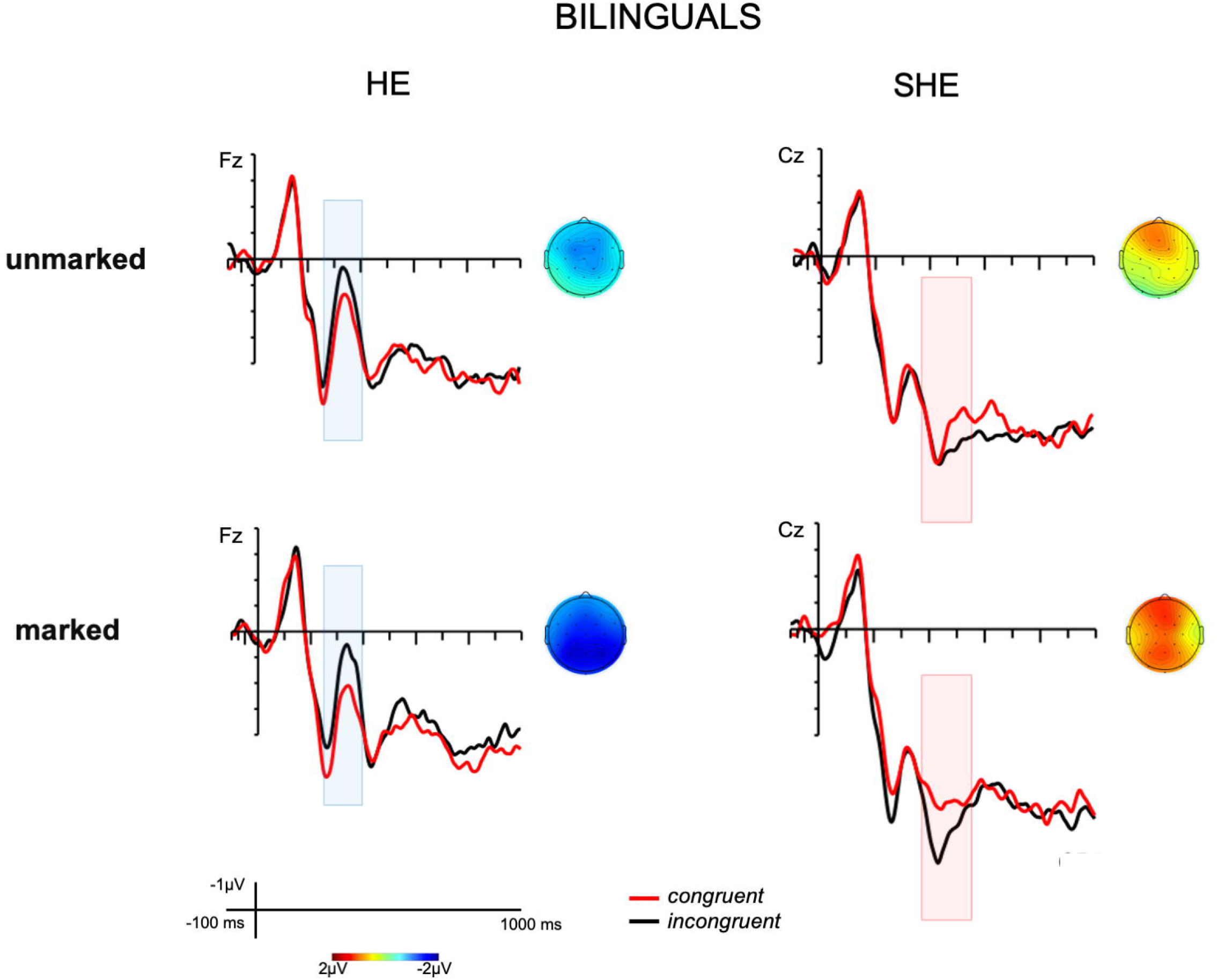
Grand-averaged ERP waveforms elicited by target pronouns in the monolingual group (upper panel) and bilingual group (lower panel), shown separately for the unmarked and marked conditions as a function of prime–target stereotypical congruency: congruent vs. incongruent. Negative voltages are plotted upward. Topographical maps show the scalp distribution of the mean incongruent-minus-congruent difference for each time window and condition in which a statistically significant effect was observed (*p* < .05).

### N400

The omnibus ANOVA on the N400 component showed a significant main effect of Congruency, F(1, 54) = 12.81, p < .001, η^2^p = .19, with more negative waveforms for stereotypically incongruent than congruent conditions. A significant Target Gender x Congruency interaction also emerged, F(1, 54) = 16.93, p < .001, η^2^p = .24, indicating a priming effect (larger N400 for incongruent vs congruent primes) that was present only for male target pronouns, not for female target pronouns, regardless of Group, highlighting an asymmetry found in our previous works. Additional significant interactions were observed for Target Gender x Congruency x Group, F(1, 54) = 4.36, p = .042, η^2^p = .08; Congruency x Longitude x Group, F(1.29, 69.67) = 7.47, p = .004, η^2^p = .12; and Target Gender x Congruency x Longitude x Group, F(1.18, 63.83) = 4.80, p = .026, η^2^p = .08. To further examine these three- and four-way interactions, follow-up analyses were conducted separately for each group (monolinguals vs. bilinguals) and for each target gender (female vs. male).

### Monolinguals

For male target pronouns, the ANOVA revealed a main effect of Congruency F(1, 27) = 10.21, p = .004, η^2^p = .27, with incongruent trials eliciting more negative responses than congruent trials. For female target pronouns, the ANOVA revealed a Congruency x Longitude interaction, F(1.36, 36.82) = 7.32, p = .006, η^2^p = .21; however, post hoc analyses indicated no significant priming effects (ps > .1). Together, these findings highlight a gender-stereotype asymmetry.

### Bilinguals

For male target pronouns, the ANOVA revealed a main effect of Congruency, F(1, 27) = 25.09, p < .001, η^2^p = .48, with incongruent trials eliciting more negative responses than congruent trials. In addition, the data revealed a Prime Type x Congruency x Longitude interaction, F(1.19, 32.15) = 3.97, p = .048, η^2^p = .13. Post hoc analyses showed that priming effects were significant only for the marked condition and not for the unmarked condition, and only at centro-parietal sites: central (t(27) = 4.93, p = .002) and parietal (t(27) = 6.56, p < .001), indicating that the N400 priming effects were driven by the marked condition. This interaction confirms our hypothesis that bilingual participants are influenced by the grammatical gender marking of the Spanish translation of the English words.

For female target pronouns, the ANOVA revealed a Congruency x Longitude interaction, F(1.26, 33.90) = 9.26, p = .003, η^2^p = .26; post hoc analyses indicated a priming effect only at anterior sites and in the opposite direction, with a larger N400 for congruent than incongruent trials (t(27) = 3.53, p = .012). Together, these findings support the presence of a gender-stereotype asymmetry in bilinguals as well.

### P300

The omnibus ANOVA on the P300 component revealed a significant main effect of Congruency, F(1, 54) = 10.74, p = .002, η^2^p = .17. Specifically, P300 amplitudes were larger for stereotypically incongruent than for stereotypically congruent conditions. A significant Target Gender x Congruency interaction also emerged, F(1, 54) = 12.70, p < .001, η^2^p = .19. Post hoc analyses indicated that the priming effect (larger P300 for incongruent vs congruent primes) was present only for female target pronouns (t(54) = 4.36, p < .001), and not for male target pronouns, regardless of Group. No interaction with Group was observed; therefore, no further follow-up analyses were conducted separately for monolingual and bilingual participants. Notably, the present findings extend the previously observed P300 gender stereotype asymmetry, which had so far been reported only in the word-face priming paradigm.

## Discussion

Does grammatical gender in a bilingual’s first language influence the online processing of gender stereotypes in a second language? The present study addressed this question by examining whether Spanish-English bilinguals process English gender-stereotyped role nouns differently depending on whether their L1 translation equivalents are gender-marked or unmarked. Using an ERP noun-pronoun priming paradigm similar to that used by Pesciarelli et al. (2019), we tested whether neural responses to English pronouns varied as a function of stereotype congruency and of the grammatical-gender marking of the primes’ Spanish translation equivalents.

The results revealed two main findings. First, both Spanish-English bilinguals and English monolinguals showed evidence of gender stereotype priming in the N400 and P300 components. Crucially, in bilinguals, the N400 effect was driven by the gender-marked condition: a reliable N400 priming effect emerged for incongruent male pronouns when English primes had gender-marked Spanish translation equivalents, but not when they had unmarked translations. This marked/unmarked modulation was specific to male pronouns and did not extend to the P300. Importantly, no comparable marked/unmarked modulation was observed in monolingual controls. Second, the stereotype priming effect showed an asymmetric ERP pattern as a function of the direction of the violation, regardless of group, with male pronouns following female-stereotyped nouns eliciting a larger N400 than congruent continuations, and female pronouns following male-stereotyped nouns eliciting a larger P300.

The first finding provides evidence consistent with our hypothesis that grammatical gender in a bilingual’s first language can influence gender stereotype processing in a second language. Specifically, it suggests that grammatical gender information from Spanish may have contributed to the online processing of English pronouns, even though gender was not overtly marked in the English nouns. Because relevant psycholinguistic variables were controlled and no comparable modulation was observed in monolingual controls, the effect is unlikely to reflect item-level differences in the English materials and is instead consistent with bilingual cross-linguistic influence from the non-target language. This finding extends previous evidence that bilinguals may activate non-target language representations even when performing a task entirely in the target language (Kroll & Stewart, 1994; Sunderman & Kroll, 2006; Thierry & Wu, 2007). Notably, the grammatical-gender translation modulation emerged for the N400 but not for the P300, suggesting that L1 grammatical gender may have primarily affected early expectancy-based integration rather than later attentional or updating processes. In other words, Spanish gender marking may have strengthened the gender expectation generated by the prime when the subsequent pronoun was processed as an early semantic mismatch. Because this effect emerged specifically in the N400 and for male pronouns following female-stereotyped nouns, it may be connected to the broader asymmetry in ERP responses to female- and male-stereotyped roles, an issue we return to below when discussing the asymmetric directionality of the ERP effects.

This bilingual-specific marked/unmarked modulation can be interpreted in light of two major models of bilingual lexical processing: the Revised Hierarchical Model and the Bilingual Interactive Activation Plus model, or BIA+. According to the Revised Hierarchical Model, bilingual lexical representations are linked across languages and to a shared conceptual system, with the strength of these links depending on dominance, proficiency, and language experience (Kroll & Stewart, 1994). In the present study, this framework helps explain why English L2 nouns may have activated their Spanish L1 translation equivalents, allowing grammatical-gender information from Spanish to contribute to the gender expectation generated by the English prime.

The BIA+ model provides a complementary account of how this influence may have occurred online. Because this model assumes language-nonselective lexical access, Spanish lexical representations may have been co-activated during the processing of English primes, even though the task was entirely in English (Dijkstra & van Heuven, 2002). From this perspective, the larger N400 priming effect for gender-marked than unmarked translation equivalents in bilinguals is consistent with the parallel activation of Spanish grammatical-gender properties during English comprehension.

Together, these models help account for the bilingual-specific marked/unmarked modulation: the Revised Hierarchical Model explains why L1 translation equivalents may influence L2 processing, whereas BIA+ explains how such influence may arise in real time through cross-linguistic co-activation. However, they do not by themselves explain the second finding: the gender-direction asymmetry observed regardless of group. This shared male-female asymmetry appears instead to reflect broader social-cognitive mechanisms involved in the processing of male- and female-stereotyped roles.

Specifically, the N400 effect for male pronouns suggests that these incongruent continuations were more difficult to integrate with the gender expectation generated by the preceding role noun. This interpretation is in line with the functional role of the N400, which is typically associated with semantic expectancy and integration processes, with a larger amplitude observed when incoming information is less expected or semantically incongruent with the preceding context (Kutas & Federmeier, 2011). In the present study, male pronouns following female-stereotyped primes, therefore, appear to have violated an expectation already activated by the role noun. This interpretation is consistent with previous ERP evidence showing that violations of female-stereotyped roles by male agents are often associated with N400 modulations (Proverbio et al., 2018; Serafini & Pesciarelli, 2025, 2026; Siyanova-Chanturia et al., 2012). More generally, this pattern fits with the idea that female-stereotyped roles may generate relatively strong and more specific gender expectations, making male pronouns immediately incongruent at the semantic level. This may also explain why the marked/unmarked modulation was restricted to the N400 effect for male pronouns: Spanish grammatical-gender marking may have strengthened these expectations, increasing the semantic mismatch when a male pronoun followed. By contrast, female pronouns following male-stereotyped primes appeared to engage a later evaluative stage, which may have been less sensitive to L1 grammatical-gender information.

By contrast, the P300 effect for female pronouns is especially noteworthy because, to our knowledge, this is the first evidence of a P300 modulation for gender stereotype violations in a purely word-pronoun priming paradigm. Previous studies have reported later positive effects for violations of male-stereotyped roles by female agents, including P300, P600, and LPP effects (Proverbio et al., 2017; Su et al., 2016; Serafini & Pesciarelli, 2025, 2026). However, P300 modulations in response to gender stereotype violations have mainly been observed in mixed word-face priming paradigms, particularly when stereotype-incongruent faces followed role nouns (Rodríguez-Gómez et al., 2020; Serafini & Pesciarelli, 2025, 2026). By contrast, previous word-pronoun studies have mainly reported N400 effects for stereotype-incongruent male pronouns, with no reliable P300 effects for stereotype-incongruent female pronouns (Siyanova-Chanturia et al., 2012; Pesciarelli et al., 2019). Thus, the present study extends previous evidence by showing that later attentional or context-updating processes may also be engaged by stereotype-incongruent pronouns, even when the target is purely linguistic. Functionally, this P300 effect suggests that incongruent female pronouns engaged a later process, possibly related to attention allocation, salience detection, or context updating. The P300 has often been associated with context updating and the evaluation of task-relevant or salient stimuli (Donchin, 1981). In the present paradigm, female pronouns following male-stereotyped primes may have been treated less as an early semantic expectancy violation and more as a salient mismatch requiring updating of the representation generated by the prime. One possible reason for the emergence of this effect is that previous word-pronoun studies included both stereotypical and grammatical gender conditions (Siyanova-Chanturia et al., 2012; Pesciarelli et al., 2019), whereas the present study focused only on stereotype congruency. In those studies, a substantial proportion of the stimuli did not involve stereotype-based expectations, but rather grammatical gender information. In the present study, by contrast, the critical manipulation was entirely based on the congruency between the stereotypical gender of the prime and the gender of the target pronoun. This may have made the stereotype dimension relatively more salient, despite the inclusion of filler stimuli, increasing participants’ sensitivity to the stereotypical gender of the primes, allowing later updating processes to emerge even with pronoun targets.

## Conclusion

Overall, the present findings provide initial evidence that grammatical gender information from a bilingual’s L1 may modulate online gender-stereotype processing in the L2. Rather than being driven only by social-conceptual expectations associated with English role nouns, bilingual stereotype processing appears to reflect an interaction between expectations generated in the target language and grammatical information activated from the non-target language. This finding represents an important first step in showing that cross-linguistic activation can shape the processing of social stereotypes, opening new avenues for examining how language-specific structures interact with social cognition during comprehension. Because this is an initial investigation, future work will be needed to assess the generalizability of these findings across different bilingual contexts, languages, and tasks.

