## Supplementary materials for "When Grammatical Gender Shapes Gender Stereotypes: Neural Evidence for Cross-Linguistic Modulation in Spanish-English Bilinguals"

This file includes:
Supplementary Methods
List of Experimental stimuli
Table 1

**Prime-word norming study**

A total of 318 English words whose Spanish translations were grammatically gender-marked and 264 English words whose Spanish translations were gender-unmarked were divided into two separate questionnaires. Corresponding Spanish versions were created using the translations of the same words. The English versions were administered to native English speakers, whereas the Spanish versions were administered to native Spanish speakers. For each language, each questionnaire was completed by 20 participants.

For the stereotypicality ratings, participants indicated the extent to which each word was associated with men, women, or both on a seven-point Likert scale (1 = only men, 4 = both men and women, 7 = only women). The same participants also completed a separate valence questionnaire, using a seven-point Likert scale (1 = negative, 4 = neutral, 7 = positive). The same two word sets were used across the stereotypicality and valence questionnaires. However, to prevent repeated exposure to the same items, participants who rated the gender-marked words for stereotypicality rated the gender-unmarked words for valence and vice versa. None of these participants took part in the main experiment.

A total of 156 English words that received high stereotypicality ratings in both the English and Spanish questionnaires were selected as experimental primes. The final set comprised 39 female- and 39 male-stereotyped English words with gender-marked Spanish translations, and 39 female- and 39 male-stereotyped English words with gender-unmarked Spanish translations. In addition, 40 English words rated as gender-neutral in both languages were selected as fillers, including 20 words with gender-marked Spanish translations and 20 with gender-unmarked Spanish translations.

### List of experimental stimuli

| MARKED | | | UNMARKED | | |
|---|---|---|---|---|---|
| MASCULINE | FEMININE | FILLER | MASCULINE | FEMININE | FILLER |
| ARCHITECT | AFFECTIONATE | AUTHOR | AGENT | ASSISTANT | ABUNDANT |
| ASTRONOMER | BABYSITTER | CITIZEN | ASSAILANT | CHEERFUL | COMPETENT |
| AUTHORITARIAN | BEAUTIFUL | CONSULTANT | BODYGUARD | DOCILE | CONFIDANT |
| BARBER | BOTANIST | CREATOR | BRICKLAYER | ELEGANT | EXHILARATING |
| BEGGAR | BUSTY | DIRECT | CHILDISH | ELOQUENT | FIERCE |
| BLACKSMITH | CALM | EMPLOYEE | COMBATANT | EMOTIONAL | ILLUSTRIOUS |
| BOSS | CAREFUL | FOLLOWER | COMEDIAN | ENDEARING | IMPOTENT |
| BOXER | CAREGIVER | GUEST | COMMANDER | EXTRAVAGANT | INDECENT |
| BUILDER | CARING | HEARTBREAKING | CONDESCENDING | FAITHFUL | INDULGENT |
| CAPTAIN | CHARITABLE | HOST | DELINQUENT | FERTILE | PLEADING |
| CARELESS | COMMUNICATIVE | INSTIGATOR | DEMANDING | FLOURISHING | PROFESSIONAL |
| CARPENTER | COMPASSIONATE | PARTNER | DOMINANT | FRAIL | RESISTANT |
| DARK | DANCER | PITIFUL | FAST | FRIENDLY | SKETCHER |
| ENGINEER | DELICATE | PRESUMPTUOUS | FORESTER | GENTLE | SKILLFUL |
| FARMER | DESIGNER | RADIOLOGIST | GALLANT | HELPER | STABLE |
| FIREFIGHTER | DRAMATIC | SHOPKEEPER | HOSTILE | INNOCENT | SURPRISING |
| FISHER | EDUCATOR | SHREWD | IMPASSIVE | LISTENER | TIRELESS |
| LIAR | ENCHANTING | SILENT | IMPERTINENT | LOVER | TRAVELER |
| LIEUTENANT | GENEROUS | SPEAKER | INSENSITIVE | LOYAL | UNREPEATABLE |
| LOCKSMITH | HAIRDRESSER | VAIN | JANITOR | NICE | WALKER |
| MECHANIC | INTUITIVE | | KEEPER | OBEDIENT | |
| MINISTER | KINDHEARTED | | LEADER | OBLIGING | |
| PHILOSOPHER | LAUNDRESS | | MANUFACTURER | PATIENT | |
| PLUMBER | LIBRARIAN | | MERCHANT | PLEASANT | |
| POLITICIAN | LOVING | | MILITANT | SENSITIVE | |
| PRODUCER | NURSE | | MILITARY | SINGER | |
| PROGRAMMER | PRECIOUS | | NAVIGATOR | SMILING | |
| PROSECUTOR | PSYCHOLOGIST | | OFFICIAL | SPARKLING | |
| RAPPER | SECRETARY | | PARAMILITARY | SPICY | |
| REFEREE | SEDUCTIVE | | RIDER | SPIRITUAL | |
| SERGEANT | SERENE | | SMELLY | STUNNING | |
| SHEPHERD | SHOPPER | | STRONG | SWEET | |
| SHOEMAKER | SLIM | | TITLEHOLDER | TALKATIVE | |
| SKATER | SUBMISSIVE | | TRADER | THEATRICAL | |
| SMOKER | SUPPORTIVE | | UNFAITHFUL | UNFORGETTABLE | |
| SURGEON | TEACHER | | UNPLEASANT | VIRGIN | |
| THIEF | TEARFUL | | UNSETTLING | VIVACIOUS | |
| TRAITOR | UNDERSTANDING | | VILE | VOCALIST | |
| VAGABOND | WITCH | | WEALTHY | WEAK | |

**Table 1**
Mean log frequency, length, stereotypicality, valence, and cognate status for marked and unmarked gender stimuli.

| | **Marked** | | **Unmarked** | | |
|---|---|---|---|---|---|
| | **feminine** | **masculine** | **feminine** | **masculine** | |
| **Log frequency** | 0.93 (0.0-2.2) | 0.88 (0.1-2.4) | 1.04 (0.3-2.1) | 0.89 (0.0-2.2) | P > .05 |
| **Length** | 8.44 (4-13) | 7.72 (4-13) | 7.59 (4-13) | 8.28 (4-13) | P > .05 |
| **Stereotypicality (raw ratings)** | 5.68 (5.0-6.6) | 2.32 (1.4-3.0) | 5.61 (5.0-6.7) | 2.46 (1.2-3.0) | P > .05 |
| **Stereotypicality (normalized)** | 2.32 (1.4-3.0) | 2.32 (1.4-3.0) | 2.39 (1.3-3.0) | 2.46 (1.2-3.0) | P > .05 |
| **Valence** | 2.55 (1.3-6.0) | 3.49 (1.6-6.5) | 2.63 (1.4-6.1) | 4.32 (2.0-6.4) | P < .05* |
| **Cognates** | 5.21 (1.0-10) | 5.10 (1.0-10) | 4.36 (1.0-10) | 4.67 (1.0-10) | P > .05 |

Min–max ratings are indicated in parentheses. T-tests were performed on the means. Stimuli were closely matched for the above properties, *except for valence, which differed significantly in all comparisons except between marked and unmarked feminine stimuli.
